# SARS-CoV-2 structure and replication characterized by *in situ* cryo-electron tomography

**DOI:** 10.1101/2020.06.23.167064

**Authors:** Steffen Klein, Mirko Cortese, Sophie L. Winter, Moritz Wachsmuth-Melm, Christopher J. Neufeldt, Berati Cerikan, Megan L. Stanifer, Steeve Boulant, Ralf Bartenschlager, Petr Chlanda

## Abstract

Severe acute respiratory syndrome coronavirus 2 (SARS-CoV-2), the causative agent of the COVID19 pandemic, is a highly pathogenic β-coronavirus. As other coronaviruses, SARS-CoV-2 is enveloped, replicates in the cytoplasm and assembles at intracellular membranes. Here, we structurally characterize the viral replication compartment and report critical insights into the budding mechanism of the virus, and the structure of extracellular virions close to their native state by *in situ* cryo-electron tomography and subtomogram averaging. We directly visualized RNA filaments inside the double membrane vesicles, compartments associated with viral replication. The RNA filaments show a diameter consistent with double-stranded RNA and frequent branching likely representing RNA secondary structures. We found that assembled S trimers in lumenal cisternae do not alone induce membrane bending but laterally reorganize on the envelope during virion assembly. The viral ribonucleoprotein complexes (vRNPs) are accumulated at the curved membrane characteristic for budding sites suggesting that vRNP recruitment is enhanced by membrane curvature. Subtomogram averaging shows that vRNPs are distinct cylindrical assemblies. We propose that the genome is packaged around multiple separate vRNP complexes, thereby allowing incorporation of the unusually large coronavirus genome into the virion while maintaining high steric flexibility between the vRNPs.

---

*Coronaviridae* is a large family of single-stranded positive-sense (+)RNA viruses that infect a broad range of vertebrate hosts. β-coronaviruses, including SARS-CoV-1 and Middle Eastern Respiratory Syndrome Coronavirus (MERS-CoV) are highly contagious pathogens that can cause severe lower respiratory infections. At the end of 2019, SARS-CoV-2 emerged in the city of Wuhan, China, likely through zoonotic transmission via a bat reservoir and a still unidentified intermediate host that subsequently led to a pandemic^3^, affecting over 17 million individuals and causing close to 700,000 deaths worldwide^5^.

Cryo-electron microscopy studies of SARS-CoV-1 and the closely related murine hepatitis virus (MHV) show that the virions are predominantly spherical or ellipsoidal with an average envelope diameter between 80 and 90 nm^6^. The main structural components of coronaviruses are the glycoprotein S, the transmembrane proteins M and E and the nucleoprotein N, which forms a viral ribonucleoprotein (vRNP) complex with the viral RNA (vRNA). SARS-CoV-2 and SARS-CoV-1 S are structurally similar glycosylated homotrimers^2,7^ that bind to the angiotensin-converting enzyme 2 (ACE2) receptor present on the cell surface of permissive cells^8^. Similar to other (+)RNA viruses^9^, coronaviruses modify cellular membranes to form double-membrane vesicles (DMVs), which are used as dedicated sites for vRNA replication^10^. Several studies of SARS-CoV-1 and MERS-CoV point towards DMVs being derived from ER cisternae in a process predominantly driven by the nonstructural protein 3 (nsp3) and nsp4^11^. In this model, the ER lumen constitutes the space between the DMV’s inner and outer membrane, while the enclosed space is of cytoplasmic origin and enriched in double-stranded RNA (dsRNA)^12^. A recent study by Wolff *et al.*^13^ identified a molecular pore complex which interconnects the DMV interior with the cytoplasm, possibly allowing RNA import and export. SARS-CoV-1 M, E and N are required for virion assembly^14^ that takes place on the cytoplasmic side of the ER-Golgi intermediate compartment (ERGIC) cisternae^15,16^. M and E proteins act as an assembly hub, interacting with both N and S proteins, thus ensuring vRNP incorporation into the nascent virion^17,18^.

Here, we used cryo-electron tomography (ET) on cryo-focused ion beam (cryo-FIB) milled lamellae or whole cell cryo-ET of various SARS-CoV-2 infected cell lines, which were chemically fixed for biosafety reasons, to structurally characterize DMV morphology, virus assembly, and extracellular virions close to their native state. In addition to commonly used monkey kidney derived VeroE6 cells, we included the human pulmonary cell lines Calu3 and A549, the latter stably expressing the ACE2 receptor (A549-ACE2), which renders these cells permissive for SARS-CoV-2 as reported for SARS-CoV-1^19^ (Fig. S1a–c). Although ACE2 mRNA levels in A549-ACE2 cells were higher than in Calu3 and VeroE6 cells, VeroE6 and Calu3 cells showed a higher proportion of infected cells at 16 hours post infection (hpi) characterized by anti-dsRNA staining (Fig. S1d, e). This indicates that ACE2 mRNA levels do not directly correlate with permissiveness of these cell lines to SARS-CoV2 infection.

## DMVs and RNA filaments

We first focused on the characterization of DMVs, which were found in all analyzed cell lines (Fig. S2), and on the visualization of RNA in the DMV interior. Cryo-ET revealed that both inner and outer DMV membranes were separated by a luminal spacing of 18 nm (SD = 9 nm, n = 32) (Fig. 1). DMV appearance and size distributions were similar in all infected cell lines with an inner average diameter of 336 nm (SD = 50 nm, n = 20) (Fig. 1c). This is in agreement with the diameter of the DMVs in SARS-CoV-1 infected VeroE6 cells, which was reported 300 ± 50 nm as measured from the outer membranes^12^. Thin and electron-dense filaments, presumably representing vRNA, were clearly observed in all DMVs (Fig. 1, Movie S1). Individual vRNA filaments appeared smooth and did not organize into bundles or concentric layers (Fig. 1d, e,) that were observed for the dsRNA of the Flock House Virus^20^. The filaments had a uniform average diameter of 2.68 nm (SD = 0.23 nm, n = 80 from two tomograms) (Fig. 1f), which corresponds well to the diameter of the A-form RNA double-stranded helix^21^ and is in accord with anti-dsRNA immuno-EM SARS-CoV-1 studies^12^. We regularly observed filament branching points (Fig. 1g, h) resembling those observed by cryo-EM studies performed on purified viral RNA^22^. However, we could neither identify an additional electron density that could be attributed to a replicase complex in the branching point nor a decrease in the diameter of the branched filaments that would indicate the presence of single-stranded RNA. This suggests that the branching points likely represent secondary RNA loops similar to those shown on purified viral RNA lacking any proteins^22^. Tracing individual RNA filaments revealed a variable length ranging from 8 – 263 nm with an average length of 52 nm (SD = 42 nm, n = 101) (Fig. 1i). This analysis likely underestimates the length of vRNA in DMVs because the cryo-lamellae are 150-200 nm thick and do not contain the entire DMV. In addition, we observed several sites where the inner and outer DMV membrane were clamped together (Fig S3), most likely corresponding to the proteinaceous pore complex reported recently^13^. Together, our observations show a DMV interior containing RNA rich in secondary structures. Although DMVs might be a site of replication, it is as well possible that its purpose is to spatially separate a ready-to-use pool of RNA transcription and replication intermediates that would otherwise be recognized by pattern recognition receptors of the innate immune system. Interestingly, the vault complexes, that among other functions have a role in dsRNA or virus-induced proinflammatory response^23^, were observed in the proximity of DMV membranes (Fig. S4a–f).

**Figure 1.**
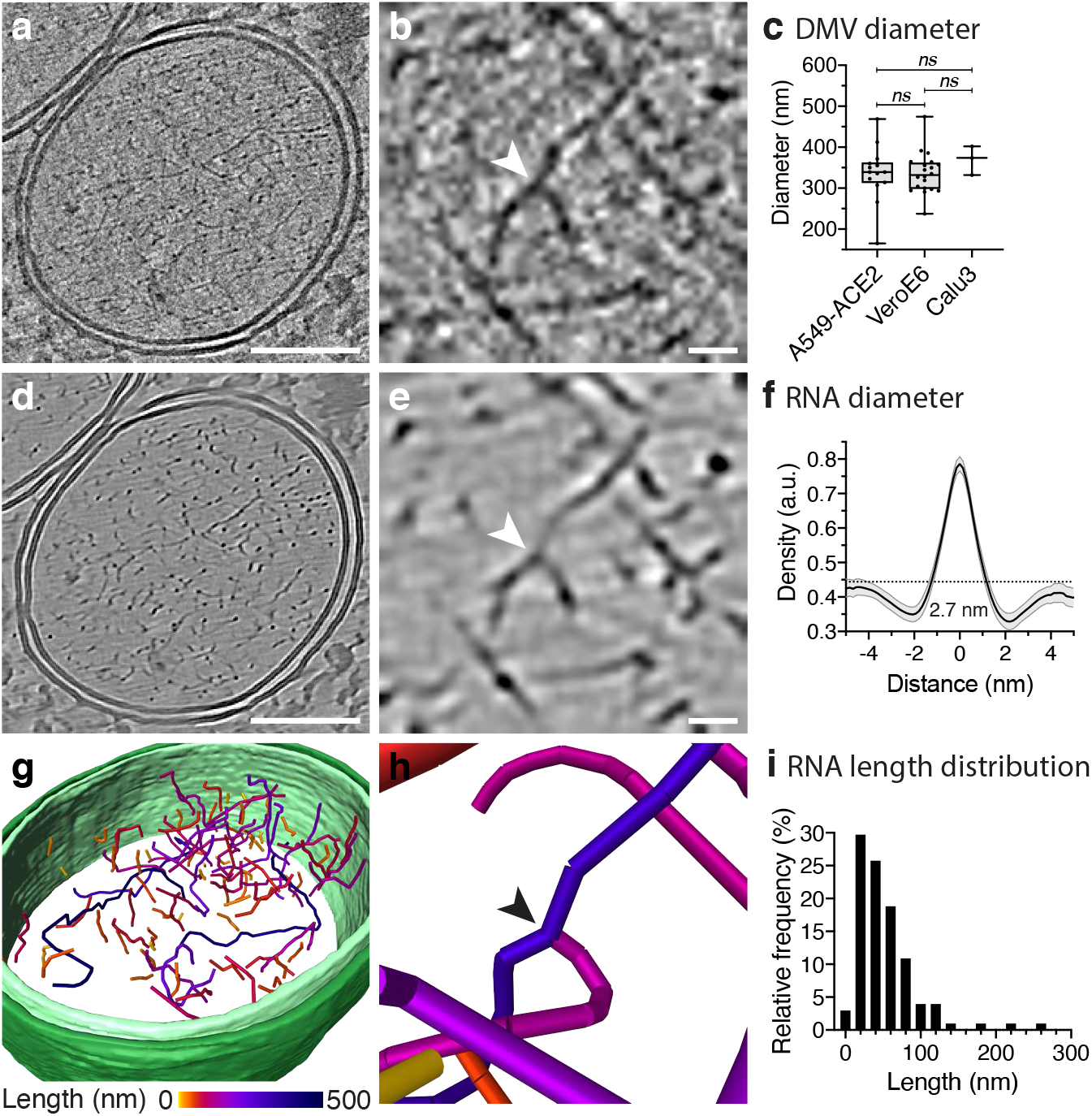
Spatial distribution and length of RNA filaments. **a**, Slice of a SIRT like filtered tomogram of an A549-ACE2 cell infected with SARS-CoV2 showing a DMV. **b**, Magnified slice of a tomogram (1.3 nm thick) showing RNA filaments and a branching point (arrowhead) in detail. **c**, Distribution of DMV inner membrane diameter in three different cell lines (A549-ACE2: n = 14; VeroE6: n = 20; Calu3: n = 3). Data is shown as Box and Whiskers plots indicating the median, 25% and 75% quantiles, minimum and maximum values and all data points. **d, e**, Tomogram slices shown in (a) and (d) after content-aware denoising using cryo-CARE1. **f**, Average normalized density line profile of filament cross-sections with indicated standard deviation (n = 80 from two tomograms) and average grey value of the DMV interior (0.44 a.u., dotted line). **g, h**, Manual segmentation of the denoised DMV (Movie S1), inner and outer membrane are represented in light and dark green, respectively. Individual segmented filaments are colored according to their length and a branching point is indicated by an arrowhead in the magnified segmentation image (h). **i**, Histogram of RNA length (n = 101) with a bin size of 20 nm. Branched filaments were measured individually. Bicubic interpolation was applied for image smoothing in (b) and (e). Scale bars: (a, d) 100 nm, (b, e) 10 nm.

It has previously been shown that DMVs are part of a network and can fuse into multivesicular compartments^12^.

To provide further structural information on how DMV fusion is mechanistically governed, DMVs within close proximity to each other were analyzed (Fig. 2). Besides funnel-like connections (Fig. 2a), we observed tightly apposed DMVs where all four membranes were stacked, forming a curved junction budding into juxtaposed DMVs (Fig. 2b, c). The junction appeared to be electron-dense, yet no particular features were found between the individual membranes. The membrane stack at the center of the junction had an average thickness of 19.4 nm (SD = 2.7 nm, n = 4), approximately conforming to four lipid bilayers. Consistent with a recent study, we also found vesicle packets (VPs) containing two or more vesicles surrounded by one outer membrane, presumably a product of DMV-DMV outer membrane fusion (Fig. 2d–f)^24^. In one case, we observed that the inner membranes of the DMVs formed a junction that resulted in an opening between two fusing inner vesicles (Fig. 2f). Based on these observations, we propose that DMV homotypic fusion occurs through membrane stacking engaging predominantly the outer, and less frequently also the inner membranes. Tight membrane apposition together with high curvature might energetically contribute to membrane fusion or to a membrane rupture followed by membrane resealing. Since DMV fusion leads to a minimization of membrane surface to volume ratio and the number of VPs increases during the infection^24^, we speculate that DMV fusion is required for repurposing membranes for virion budding at advanced stages of the replication cycle, which is supported by the observation of a fully assembled virion inside the VP lumen (Fig. 2d).

**Figure 2.**
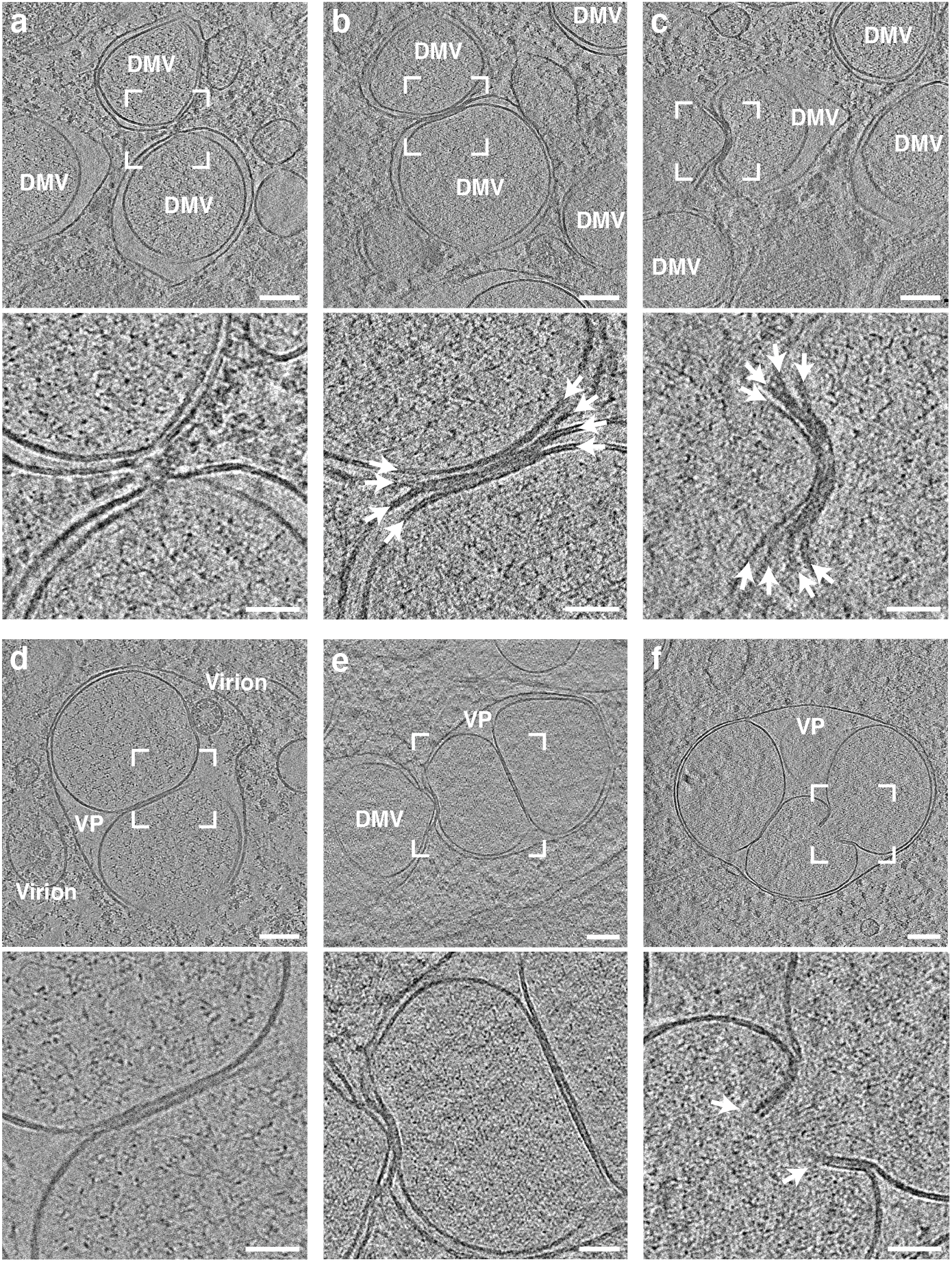
Juxtaposed DMVs form membrane junctions that lead to homotypicfusion. Tomograms showing DMVs in VeroE6 (a-c) and Calu3 (d-f) cells at 16 hpi. For each tomogram, a magnified area is shown below and indicated as a white square. 10 slices of the tomogram were averaged. **a**, DMV-DMV interaction via a constricted outer membrane connection. **b, c**, Tightly apposed membrane stack composed of four membranes (arrows) of varying curvature between two juxtaposed DMVs. **d**, VP containing two inner vesicles with a tight membrane membrane contact. **e**, DMV interaction with a VP containing two inner vesicles. **f**, VPs containing four inner vesicles with tight contact and an opening aperture formed by two stacked membranes indicated by white arrows. Scale bars: (a-f) 100nm, (magnified areas) 50 nm.

## Virion assembly and structure of intracellular virions

The ERGIC constitutes the main assembly site of coronaviruses^15^ and budding events have been described by EM studies^12,25,26^. *In situ* cryo-ET allowed us here for the first time to study SARS-CoV-2 virion assembly in close to native conditions and enabled us to localize individual S trimers and vRNPs with high precision. Virus-budding was mainly clustered in regions with a high vesicle density and close to ER- and Golgi-like membrane arrangements (Fig. 3a, S5, Movies S2, S3). S trimers were regularly found in low quantities with the ectodomain facing the ERGIC lumen. Even at high concentrations, in the absence of vRNPs, S trimer accumulation did not coincide with positive membrane curvature (Fig. 3b, c), indicating that S alone is unable to initiate virus budding. This is consistent with a study showing that SARS-CoV-1 virus-like particles are formed and released in the absence of S^14^. Early budding events with a positively curved membrane were decorated on the lumenal side with S and on the cytosolic side with vRNPs as separate cylindrical complexes (Fig. 3d, e) in all observed events (n = 19). This suggests that vRNPs may contribute to membrane curvature after recruitment to the membrane by the M protein or they might accumulate preferentially at curved membranes. Unlike matrix proteins of other enveloped viruses (e.g. influenza A, Ebola virus), our data show that M does not form a clearly discernible matrix layer and thus might alone not be able to curve the membrane. This is consistent with previous data on SARS-CoV-1 showing that expression of M alone is not sufficient to induce virion assembly but requires E and N^14^, while the assembly of other coronaviruses such as MHV is N independent^27^. Budded virions that were located directly adjacent to the ERGIC membrane showed a polarized distribution of S trimers facing towards the ERGIC lumen (Fig. 3f), likely due to steric hindrance. In contrast, virions that were more distant from the membrane showed a dispersed distribution of S (Fig. 3g) around the entire virion, indicating that S trimers are mobile on the virion envelope and redistribute during the budding process. Thus, the previously proposed lattice between S – M – N^28^ is likely not rigid and the S protein may laterally move.

**Figure 3.**
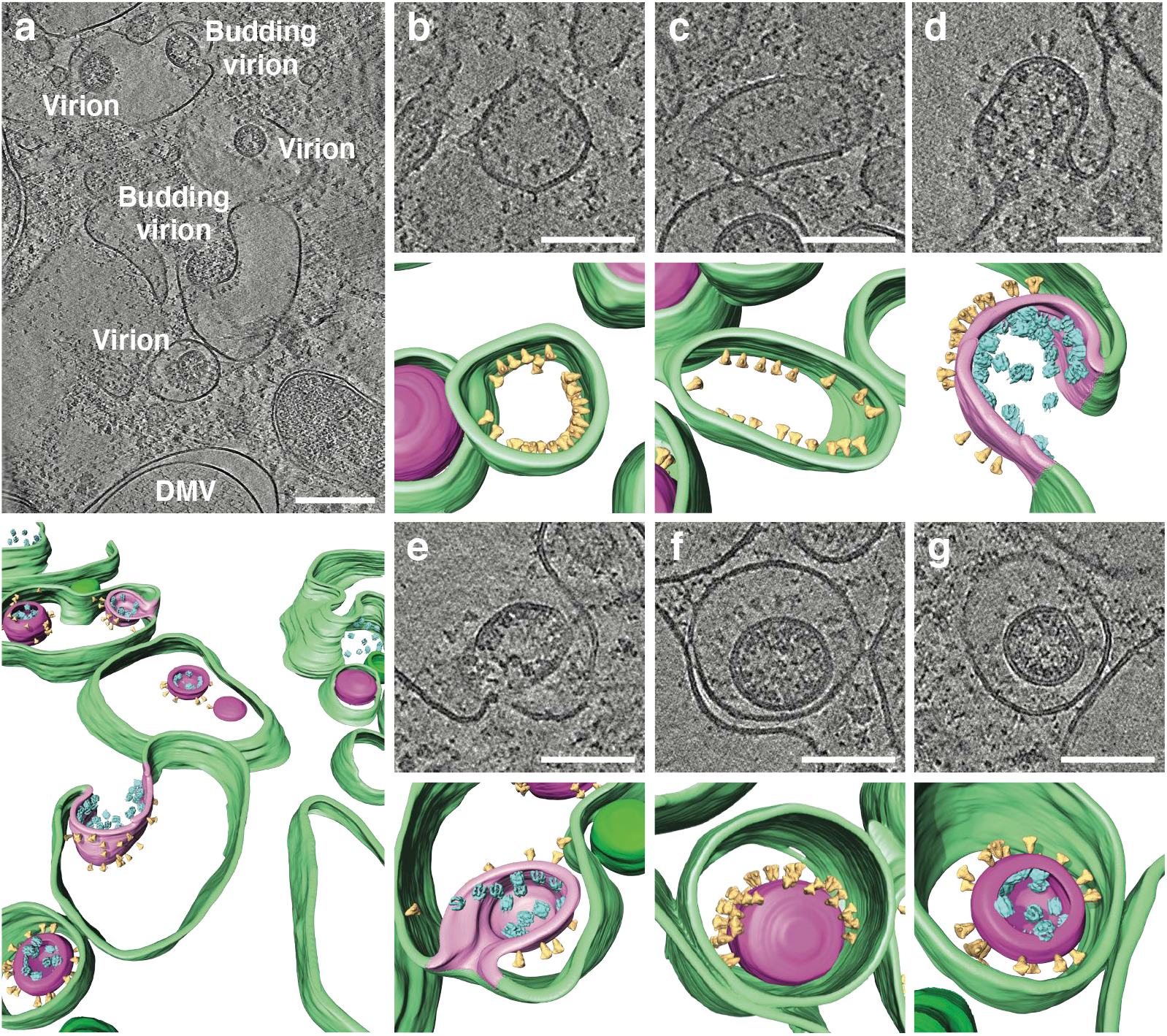
SARS-CoV-2 virion budding and assembly at the ERGIC membrane. Different budding events captured in two tomograms (Fig. S5, Movies S2, S3) of VeroE6 cells infected with SARS-CoV-2 at 16 hpi. A 3D volume rendering is shown for each area with cellular and viral membranes in green and magenta, respectively. S (yellow) and vRNPs (cyan) are represented as subtomogram averages. The S and vRNP locations correspond to the location in the tomogram, vRNP orientations were randomized. A non-local means filter was applied, and 20slices were averaged. **a**, Overview of budding events at the ERGIC membrane and intracellular released virions inside the ERGIC lumen are indicated. **b, c**, Accumulated S at the lumenal side of the ERGIC membrane. **d, e**, Early virion budding stage with S and vRNPs accumulated at the lumenal and cytosolic ERGIC membrane, respectively. **f**, Assembled viron in the ERGIC lumenin close proximity to the membrane, showing a polarized distribution of S. **g**, Assembled virion further away from the membrane with redistributed S. Scale bars: (a) 200 nm, (b-g) 100 nm.

Further analysis of the structure of intracellular virions revealed that S trimers are not always oriented orthogonally to the membrane of the spherical virions (Fig. S4). We observed tilting angles up to 41° (Fig. S4d) of S trimers on intracellular virions, supporting the observation in two recent studies^29,30^ that report that S trimers can tilt up to 60° in purified virions. Importantly, our study rules out that this tilting is caused by forces exerted on the virion during ultracentrifugation or blotting during the sample preparation, since the intracellular virions were protected during sample preparation by the cellular environment. Previous ultrastructural studies of other coronaviruses reported average virion diameters of 85 ± 5 nm for MHV^6^ and 88 ± 6 nm for SARS-CoV-1^28^. We did not observe significant differences in the average diameters of virions derived from the three cell lines (Fig. 4c) and calculated an average diameter of 89.8 nm (SD = 13.7 nm, n = 74). Subtomogram averaging (STA) of 219 individual S trimers from intracellular virions yielded a low resolution structure with a total height of approximately 25 nm measured from the virion envelope and a total width of 13 nm (Fig. 4d–f, S5). The density map featured a well-defined trimeric structure with a height of 16 nm which is in agreement with the published structure solved by single particle analysis of the purified S ectodomain in pre-fusion conformation truncated at serine residue 1147 (PDB:6VXX)^2^, indicating that the S trimer is fully formed during virion budding and that samples are well preserved despite the chemical fixation step that we had to use for biosafety reasons. The approximately 9 nm long gap between the trimeric density and the virion envelope that can be attributed to the triple-stranded coiled-coil heptad repeat 2 (HR2) was not resolved, because of the presence of three flexible hinges in the stem region as shown by two recent studies^29,30^. To estimate the number of S trimers per virion, we extracted the 3D coordinates of all identifiable S trimers on nine intracellular virions and determined the average nearest-neighbor distance to be 23.6 nm (SD = 8.1 nm, n = 100) (Fig. 4g). Based on the mathematical problem of arranging any number of points on a sphere to maximize their minimum distance, known also as the ‘Tammes Problem’^31,32^, we estimated the average number of S trimers per virion to be 48 with a range of 25 – 127 (Fig. S8). This is in line with our observation of heterogeneous S trimer densities on the surface of different virions and concurs with previously reported estimates of 50 – 100 S trimers per virion^28,33^.

**Figure 4.**
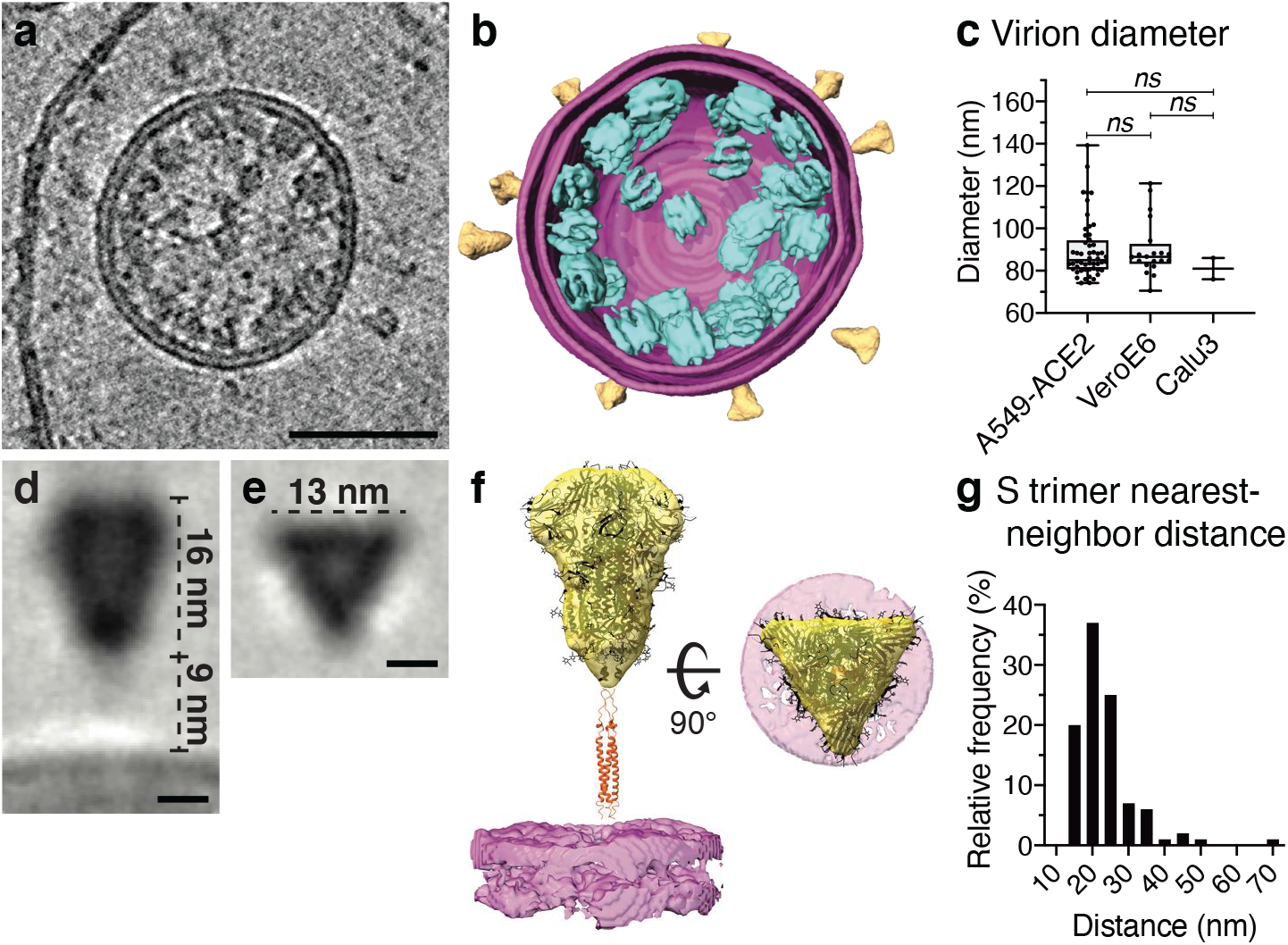
Structural analysis of intracellular virions. **a**, Tomogram showing an intracellular virion of VeroE6 cells infected with SARS-CoV-2 at 16 hpi. 20 slices of the tomogram were averaged and a median filter (radius = 1 pixel) was applied. **b**, 3Dvolume rendering of (a) with the viral envelope shown in magenta, with both leaflets of the membrane resolved. S (yellow) and vRNPs (cyan) are represented as subtomogram averages. The S and vRNP locations correspond to the location in thetomogram, vRNP orientations were randomized. **c**, Distributions of intracellular virion diameters measured in A549-ACE2 (n = 52), VeroE6 (n = 20) and Calu3 cells (n = 3). Box and Whiskers plots indicate the median, 25% and 75% quantiles and minimum and maximum values as well as all data points. Unpaired t-test showed no significant differences between the diameter of intracellular virions found in the three different cell lines. **d, e**, Central longitudinal (d) and cross-sectional slice (e) showing the subtomogram average of the S trimer of intracellular virions. **f**, Orthogonal views of the S trimer subtomogram average (yellow) and the virion envelope (magenta). Fitted structure of the Strimer ectodomain (PDB:6VXX)^2^ and the HR2 domain of SARS-CoV-1 (PDB:2FXP)^4^ are shown in black and orange, respectively (Movie S4). **g**, Plot showing the distribution of S nearest-neighbor distances (n = 100) on the surface of virions with an average of 23.6 nm (SD = 8.1 nm, n = 100). Scale bars: (a) 50 nm (d,e) 5 nm.

## Structure of extracellular virions

We next analyzed extracellular virions in the vicinity of infected cells to provide insights into conformational changes of the S trimers and vRNP structure and their arrangement within the virion. Virions were found close to the plasma membranes of all host cells, albeit with notable differences in the number of virions directly attached. We found few virions around A549-ACE2 cells, all of which appeared to directly interact with the cell surface (Fig. 5a). In contrast, a high concentration of virions from VeroE6 cells were covering the surface of filopodia and protrusions that interconnected neighboring cells (Fig. 5b, S9). Similar observations have been made in a recent study on VeroE6 and Caco-2 cells showing that SARS-CoV-2 infection induces the formation of virion decorated filopodia^34^. These observations are reminiscent of the cell-to-cell transmission via viral surfing as reported for HIV^35^ and could argue for a yet undescribed mode of transmission for SARS-CoV-2. In contrast, no virions were found attached to the surface of Calu3 cells (Fig. 5c). Differences in the number of virions attached to the analyzed cell lines might be explained by different levels of surface proteases such as TMPRSS2 or ADAM17, which cleave ACE2 receptors and thereby controlling its abundance on the cell surface^36^. In this scenario, SARS-CoV-2 virions remain attached to the cell surface after exocytosis due to S interactions with ACE2, which could control not only virion entry but also release into the surrounding environment. In agreement with our observations in intracellular virions, extracellular virions released from VeroE6 cells were studded with S trimers in pre-fusion conformation that were often tilted and had an average length of 23.4 nm (SD = 2.3 nm, n = 48) (Fig. 5e). Noticeably, extracellular virions from A549-ACE2 cells exhibited almost exclusively thin, rod-shaped S trimers resembling the post-fusion conformation^37,38^ with an overall length of 23.2 nm (SD = 2.3 nm, n = 8) and thickness of 3.8 nm (SD = 0.7 nm, n = 9) (Fig. 5d). Virions from Calu3 cells displayed a mixture of both spike conformations (Fig. 5f). The host cell-type dependent difference in the number of pre-fusion and post-fusion conformations may be explained by different levels of TMPRSS2 proteases^39^ and ACE2 receptors that trigger S conformational changes. Consistently, S trimers in post-fusion conformation were primarily detected on virions directly interacting with A549-ACE2 cells expressing high levels of ACE2 (Fig. 5d, S1). In contrast, S present on virions in ACE2 low-expressing VeroE6 cells (Fig. S1) were almost exclusively in pre-fusion conformation.

**Figure 5.**
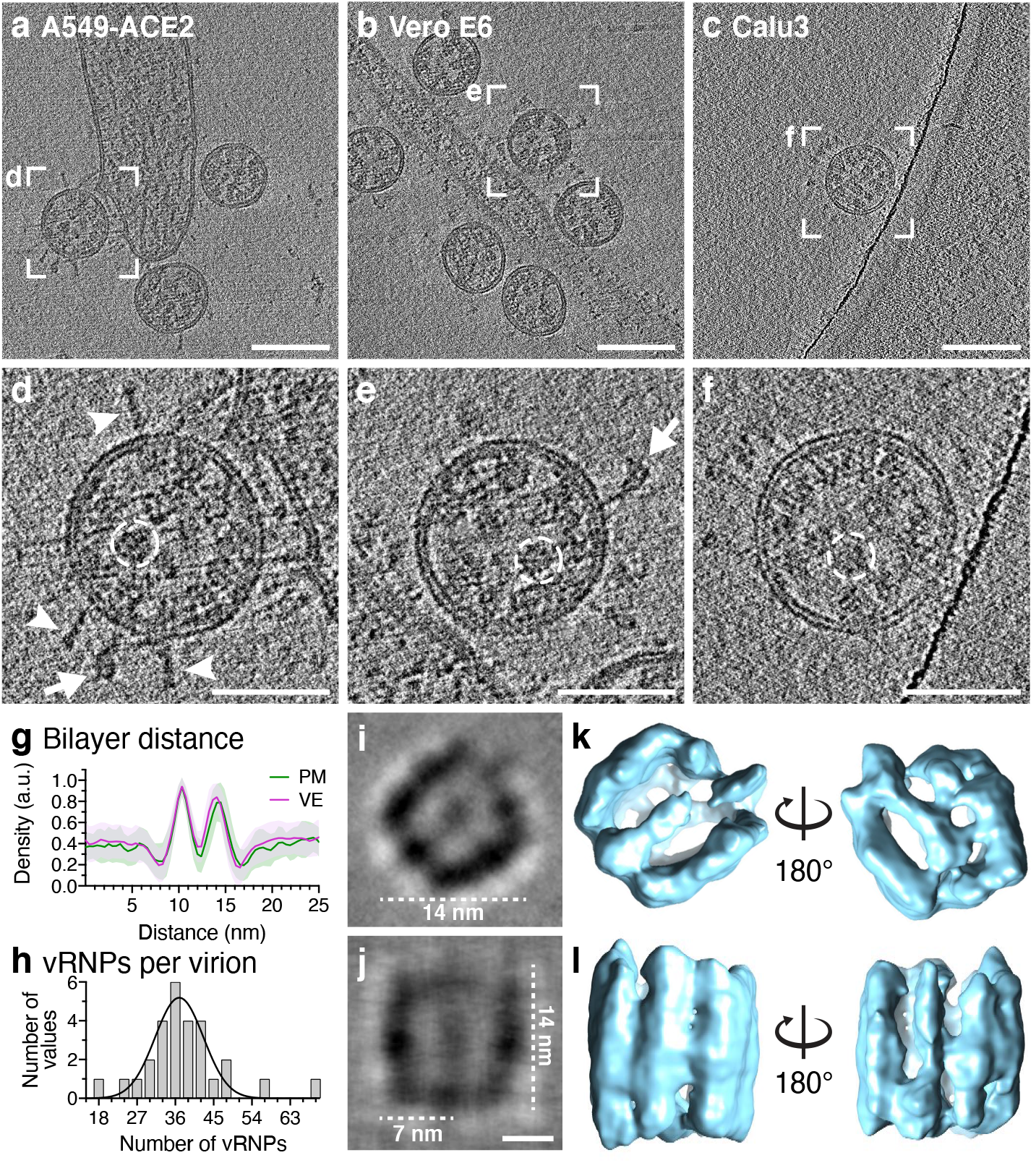
Structural analysis of extracellular virions. **a-c**, Slices of tomograms showing extracellular virions released from A549-ACE2, VeroE6 and Calu3 cells, respectively. **d-f**, magnified views of (a-c), for better visualization, 10 slices were averaged. Exemplarily, S glycoproteins in a pre-fusion (arrow) or post-fusion conformation (arrowhead) are marked, vRNP complexes are encircled with dashed lines. **g**, Plot profile through the viral envelope (VE, magenta) of virions released fromVeroE6 cells and adjacent plasma membrane (PM, green) to determine monolayer separation (n = 129 and 49, respectively, SD indicated). **h**, Number of vRNP complexes per virion (n = 28) released from VeroE6 cells. Data is represented as a histogram with a bin size of 3 and Gaussian fit (R^2^ = 0.8645). **i, j**, STA of vRNP complexes from 1570individual vRNPs found in 15 tomograms. Central XZ-slice (I) and XY-slice (j) are shown. **k, l**, Iso surface representation of the sub tomogram average (Movie S5) shown from the top (k) and side view (l). Scale bars: (a-c) 100 nm, (d-f) 50 nm, (k, l) 5 nm.

Based on a previous cryo-ET study revealing an unusual membrane thickness of MHV virions of 7 – 8 nm^6^, we measured the lipid bilayer separation of SARS-CoV-2 virions. Density line profiles across the membranes showed a phospholipid monolayer separation of 3.6 nm (SD = 0.5 nm, n = 129) (Fig. 5g) whereas the host cell plasma membrane was slightly but significantly thicker (3.9 nm, SD = 0.5 nm, n = 49) as evaluated by a two-sided t-test (p = 0.003). Thus, the 25 kDa type III transmembrane glycoprotein M, and the membrane spanning E protein do not increase the thickness of the viral envelope. The M protein C-terminal endodomain facing the virion interior is an amphipathic, approximately 100 amino acids long domain that has been predicted to be positioned along the viral membrane^40,41^. Consistent with the predicted position and the small size of this endodomain, our data does not reveal a well-discernable matrix layer underneath the viral envelope such as for example the M1 layer in influenza A viruses^42^. Extracellular virions displayed on average 38 vRNPs per virion (SD = 10, n = 28) (Fig. 5h). Individual vRNPs associated with viral envelopes were often aligned in stacks, forming filaments with a width of approximately 14 nm (Fig. 5d–f, S10), indicating a preferred stacking orientation of vRNPs. STA of 1570 vRNPs yielded a compact, cylindrical assembly of 14 nm in length and a quasi-circular base with a similar diameter (Fig. 5i–l, S11, Movie S5). The assembly is composed of parallel-stacked, pillar-shaped densities, presumably formed by multiple linearly aligned N proteins. Pillar-shaped densities form two densely-packed curved walls opposing each other and surrounding a central density (Fig. 5i–l). While the walls are connected by an additional pillar on one side, the other side appears more flexible with two pillar-like densities creating an opening. Previous EM studies of vRNPs isolated from MHV and SARS-CoV-1 virions described a helical nucleocapsid^6,43^ or a coiled vRNP that forms a membrane-proximal lattice^28^. Here, we propose that vRNPs are composed of N proteins in complex with the viral genome, which links neighboring vRNPs like ‘beads on a string’. This would allow for efficient packing of the unusually large vRNA genome into the virus particles while maintaining high steric flexibility between the individual vRNPs required during their incorporation into budding virions. Based on our data, we can clearly distinguish individual vRNPs, while it remains to be investigated how the individual N proteins and RNA are organized in the vRNPs.

## Discussion

Our report provides *in situ* cryo-ET analysis of SARS-CoV-2 and induced DMVs at high preservation levels. Direct visualization of DMVs unveils a high concentration of branched double stranded RNA filaments as the major constituents of the DMV interior. Given the lack of protein densities in the DMV interior, it is conceivable that the viral replicase complex is tightly associated with the inner DMV membrane and releases vRNA into the DMV lumen. Alternatively, the DMV interior might serve the selective accumulation of RNA intermediates that are imported into the DMV via the recently discovered pore complex^13^ to evade the antiviral innate immune response. Our data indicate that S trimers alone do not induce membrane curvature during budding into the ERGIC lumen but are remarkably motile and flexible. We report that vRNPs are individual complexes that are often associated with the membrane and likely contribute to or require membrane curvature for their incorporation into budding virions. We propose that the cylindrical assembly of vRNPs enables efficient packaging of the genome in a nucleosome-like fashion.

## Methods

### Cells and viruses

VeroE6, HEK293T and A549 cells were obtained from ATCC and were maintained in Dulbecco’s modified Eagle medium (DMEM) containing 2 mM L-glutamine, non-essential amino acids, 100 U/ml penicillin, 100 μg/ml streptomycin and 10% fetal calf serum (DMEM complete). Calu3 cells were a kind gift from Dr. Manfred Frey, Mannheim and were maintained in DMEM complete supplemented with 10 mM sodium pyruvate and a final concentration of 20% FBS. For generation of A549 cells stably expressing ACE2, lentivirus was generated by co-transfection of HEK293T cells with pWPI plasmid encoding for ACE2 and the pCMV-Gag-Pol and pMD2-VSV-G packaging plasmids (kind gifts from Dr. D. Trono, Geneva). Supernatant containing lentiviruses were harvested two days post-transfection, filtered through a 0.44 μm filter and used for transduction of A549 cells followed by selection with neomycin (0.5 mg/ml). The Bavpat1/2020 SARS-CoV-2 virus isolate (provided by Prof. Christian Drosten at the Charité in Berlin, Germany, and provided by the European Virology Archive (Ref-SKU: 026V-03883) was used. A passage 4 working stock of this isolate was generated by passaging the virus twice in VeroE6 cells.

### Sample inactivation by chemical fixation

Cells (VeroE6, Calu3, A549-ACE2) were seeded in 35 mm Petri dishes containing Quantifoil 200 mesh R2/2 Au or Ti grids at seeding density of 2×10^5^ cells/dish one day before infection. A549-ACE2, Calu3 and VeroE6 were infected at different MOI (Table S1) with SARS-CoV-2, respectively. For optimal sample preservation, all fixation steps were performed with electron microscopy grade Paraformaldehyde (PFA) (Electron Microscopy Sciences) and Glutaraldehyde (GA) (Electron Microscopy Sciences) and all fixation solutions were buffered in 0.1 M PHEM buffer at pH 7.4. Infected cells were chemically fixed at 16 hpi by addition of a double-concentrated fixative solution (8% PFA, 0.2% GA, 0.2 M PHEM buffer, pH 7.4) directly into an equal volume of growing medium for 10 min at RT. Subsequently, the medium containing fixative was removed, and cells were chemically fixed for 30 min at room temperature with 4% PFA and 0.1% GA, 0.1 M PHEM buffer, pH 7.4. For a comparison of structure preservation, a subset of samples was inactivated by using 4% PFA and 0.1% GA in 0.1 M PHEM, pH 7.4 for 30 min at RT omitting the double-concentrated fixative solution. Finally, all 35 mm dishes containing infected cells were completely submerged into a container with 100 ml of 4% PFA in 0.1 M PHEM, pH 7.4 and carried out of the BSL3 facility to be immediately plunge-frozen.

### Plaque assay

VeroE6 cells were seeded into 24 well plates at seeding density of 2.5×10^5^ cells/well. Serial dilutions of infectious supernatants were prepared and added to the cells in duplicates. Infection was performed at 37°C for 1 h followed by removal of the supernatants and overlaying with 1 ml of plaquing medium (0.8% carboxymethyl-cellulose in MEM). Cells were incubated at 37°C for three days followed by fixation in 5% formaldehyde for 1 h. Plates were submerged in a solution of 6% formaldehyde for 30 min before being carried out of the BSL3 area. Plates were washed with water and 1 ml of staining solution (1% crystal violet in 10% ethanol) was added for 15 min. Staining solution was removed, plates were rinsed again with water, plaques were counted, and titers were calculated.

### Light microscopy

Infected cells grown on glass coverslips were fixed with 4% paraformaldehyde in PBS for 30 min at room temperature. Plates were then submerged in a solution of 6% formaldehyde for 30 min before being carried out of the BSL3 area. Samples were rinsed twice in PBS, permeabilized with 0.2% Triton-X100 in PBS for 10 min, rinsed again twice with PBS and incubated for 1 h with blocking buffer containing 2% milk in PBS. Samples were incubated with primary antibodies diluted in blocking buffer for 1h at RT, washed 3 times in washing buffer (PBS, 0.02% Tween-20) and incubated with secondary antibodies for 45 min at room temperature. Coverslips were mounted on glass slides with Fluoromount-G mounting media containing DAPI.

### RT-qPCR

The NucleoSpin RNA extraction kit (Macherey-Nagel) was used to isolate total intracellular RNA according to the manufacturer’s specification. cDNA was synthesized from the total RNA using a high capacity cDNA reverse transcription (RT) kit (ThermoFisher Scientific) according to the manufacturer’s specifications. Each cDNA sample was diluted 1:15 and used directly for qPCR analysis using specific primers and the iTaq Universal SYBR green mastermix (Bio-Rad). Primers for qPCR were designed using the Primer3 software^44^: ACE2 (forward) 5’-CACGAAGGTCCTCTGCACAA-3’, ACE2 (reverse) 5’-ATGCTAGGGTCCAGGGTTCT-3, SARS-CoV-2-N (forward) 5’-GCCTCTTCTCGTTCCTCATCAC-3’, SARS-CoV-2-N (reverse) 5’-AGCAGCATCACCGCCATTG-3’, HPRT (forward) 5’-CCTGGCGTCGTGATTAGTG-3’ and HPRT (reverse) 5’-ACACCCTTTCCAAATCCTCAG-3’. To obtain the relative abundance of specific RNAs from each sample, cycle threshold (ct) values were corrected for the PCR efficiency of the specific primer set and normalized to hypoxanthine phosphoribosyltransferase 1 (HPRT) transcript levels.

### Sample processing for cryo-EM

To generate the dataset for this study, 3 independent infections of cells grown on electron microscopy grids were done for A549-ACE2 cells and one infection on VeroE6 and Calu3 cells. Detailed overview of sample processing is shown in Table S1. Tomograms containing cellular structures that were partially compromised by fixation or poor vitrification as apparent from increased local membrane undulation and increased aggregation in the cytosol were excluded from the analysis.

### Plunge freezing

Chemically fixed infected cells grown on grids were plunge-frozen in liquid ethane cooled to −183°C by liquid nitrogen using a Leica GP2 plunger. Before plunge freezing, 3 μl of 0.1 M PHEM buffer pH 7.4 was applied on a grid in the chamber set to 70% humidity and 24°C. Grids were blotted for 3 seconds using No. 1 Whatman paper and clipped into Autogrids (ThermoFisher Scientific) dedicated for cryo-FIB milling and stored in liquid nitrogen.

### Cryo-focused ion beam milling

Lamellae were prepared using a focused Gallium ion beam on an Aquilos dual-beam FIB-SEM microscope at the stage temperature around −180°C (ThermoFisher Scientific). Cells positioned in the center of the grid squares were selected for milling and the eucentric height for each of the cells was set using the routines in the MAPS software. An organo-metallic platinum protective layer was applied for 5 – 6 sec onto the cells before milling. Rough milling was performed gradually in 4 steps at a stage angle of 15-18° using a Ga^+^ beam to obtain 500 nm thick lamellae. Final milling was performed at the end of the milling session to yield lamellae with nominal thickness of 150 nm. To minimize lamella bending, a micro-expansion joints milling pattern was applied as described by Wolff *et al.^45^*. Progress was monitored by SEM imaging at 2.2-10 kV with ETD and T2 detectors.

### Cryo-electron tomography

Cryo-electron tomography was performed using the Titan Krios cryo-TEM (ThermoFisher Scientific) operated at 300kV equipped with a K3 direct electron detector and a Gatan imaging filter (Gatan) in EFTEM nanoprobe mode. Lamellae or whole cells were mapped at a magnification of 6,500× at defocus −50 μm to localize DMVs and virions. Tilt series were acquired at magnifications of 26,000×; 33,000× or 64,000× (corresponding pixel sizes at the specimen level: 3.356 Å and 2.671 Å, 1.379 Å respectively) using SerialEM ^46^. A dose-symmetric acquisition scheme^47^ with a constant electron dose of approx. 3 e^−^/Å^2^ per projection, target defocus ranging from −5 to −2.5 μm and energy filter slit at 20 eV was used covering the tilt range from 60° to −60° with 3° increments. When possible, tilt series acquired on lamellae were tilted to 6° as a starting angle to compensate for the pre-tilt of the lamella with respect to the grid. Individual projections were acquired in counting mode using dose fractionation. 10 – 20 individual frames per projections were aligned and summed on-the-fly using the SEMCCD plugin in SerialEM.

### Tomogram reconstruction and volume rendering

Tilt series (TS) were aligned and reconstructed with the IMOD software package^48^. The stack was aligned using patch tracking for TS acquired on lamellae or gold fiducial markers for TS acquired in the periphery of whole cells. After CTF correction by phase-flipping and dose-filtration, the final reconstruction was performed by weighted back-projection with a SIRT-like filter equivalent to 10 iterations. For volume rendering, the pre-aligned TS stack from IMOD was reconstructed and denoised with the cryo-CARE library using the Tomo2Tomo even-odd scheme^1^ on 4x binned data. This denoised tomogram was further processed with Amira 2019.3 (ThermoFisher Scientific) using the Membrane Enhancement Filter with a feature scale of 6.5 nm. An initial segmentation was created using the Top-hat tool. Based on this initial segmentation, membranes were fully segmented in a manual fashion.

### Subtomogram averaging

To obtain the average of the S trimer present on the surface of intracellular virions during and after budding into the ERGIC, 219 positions corresponding to the center of the S trimers on the virion surface were manually picked from 6 tomograms acquired on cryo-lamellae using the general model available in Dynamo^49^. Subsequently, subtomograms with a cubic side length of 160 voxels size were cropped from tomograms and were iteratively aligned against a reference composed of two overlapping cylinders approximately matching the S trimer and globular region diameters. C3 symmetry was applied during the alignment steps since the C3 symmetry was revealed when it was not imposed (Fig. S7A). The atomic structures of the pre-fusion SARS-CoV-2 S ectodomain (PDB:6VXX)^2^ and the HR2 domain of SARS-CoV-1 (PDB:2FXP)^4^ were automatically using the fitting tool or manually placed into the density map in ChimeraX^50^, respectively. The average of the vRNP complexes within extracellular virions was obtained from 1570 particles manually picked from 15 tomograms using the general model in Dynamo. Subtomograms with a cubic side length of 168 voxels were cropped and 200 particles were iteratively averaged using a cylindrical mask adjusted to the dimensions of the vRNP to obtain a reference template. The initial average was then used as a template to obtain a refined average using 1570 particles.

### Estimation of number of S trimers per virion

Arranging a specific number of points on a sphere with a largest possible minimum distance between any point is a mathematical problem called ‘Tammes Problem’^31^ or the 7^th^ unsolved mathematical problem listed by Steve Smale^51^. It has been solved only for a few configurations^32^, although was heuristically approximated for others^52^. Based on these estimates and considering two simplified postulates, i.e. (i) virions are spherical and (ii) S trimers are evenly distributed on the viral envelope, we can determine the number of S trimers per virion.

First, we determined the average distance of S trimers on SARS-CoV-2 virions. S trimers were manually picked in reconstructed tomograms using the IMOD software package^48^ and their 3D-coordinates were extracted. For each individual S trimer, the nearest neighboring S trimer on the same virion was identified and the Euclidean distance was calculated using the pandas and numpy software libraries^53,54^. If two S trimers were reciprocal nearest neighbors, their distance was counted only once. To estimate the mean number of S trimers per virion, we calculated the central angle for two points on a sphere (Equation 1) with the previously determined average virion diameter and S trimer distance. The error of the central angle calculation was determined by error propagation (Equation 2). By comparing the determined central angle with the estimated results for configurations of N points on a sphere by Slalone *et al.*^52^ (Fig. S6d), we can determine the total number of S trimers per virion.

**Equation 1**: Central angle with spike distance *d* and virion radius *r*.

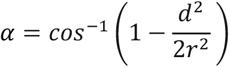

**Equation 2**: Error of central angle with spike distance error *d*_err_ and virion radius error *r*_err_

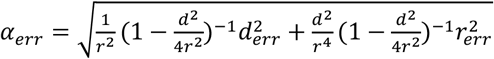

### Measurement of phospholipid bilayer distance of membranes

Membrane thickness of the virion envelope and cellular membranes was measured as the distance between phospholipid head groups of the bilayer. Line density profiles (10 pixels in width, 0.41 Å/pixel) of approximately 20 nm in length were determined across the respective membrane using the plot profile tool in FIJI/ImageJ^55^. For each membrane, 49 – 129 measurements were performed and normalized against the maxima of each measurement separately. Densities were aligned to the first maxima and plotted against the distance from the start of the line profile. The distances between the two global maxima corresponding to the phospholipid bilayer were then measured for each plot profile.

### RNA segmentation and measurement of thickness

Tracing was done manually in Amira 2019.3 using the filament editor in linear tracing mode. The reconstructed tomogram was 4x binned and either SIRT-like filtered equivalent to 10 iterations or denoised with cryo-CARE as described above. The total filament lengths were extracted from the resulting spatial graph statistics. The RNA filament diameter was measured on the unbinned SIRT-like filtered reconstructed tomogram in FIJI by smoothing the image using gaussian blur (σ = 2 px) and measuring the line profiles perpendicular to RNA filaments using FIJI’s plot profile tool (10 pixels in width, 1.379 Å/pixel). The grey value profiles were normalized, aligned to their respective maxima and averaged. The diameter was defined as the distance between the two centermost intersection points of the normalized line profile with the line corresponding to the average grey values of the DMV internal space.

## Acknowledgments

We thank Dr. Manfred Frey, Mannheim for kindly providing Calu3 cell line, Dr. Christian Drosten and the European Virus Archive-Global for providing the SARS-CoV-2 seed virus (strain Bavpat1/2020), and Christian Willig for help with the error propagation calculation. We thank Dr. Ivonne Morales and Dimitris Papagiannidis for critical reading of the manuscript. We would like to acknowledge microscopy support from the Infectious Diseases Imaging Platform (IDIP) at the Center for Integrative Infectious Disease Research Heidelberg. We would like to acknowledge access to the infrastructure and support provided by the Cryo-EM Network at the Heidelberg University (HD-cryoNet).

## Author contributions

Sample preparation: S.K., M.C., S.L.W., M.W-M., C.N., B.C., M.L.S., S.B., P.C; study conceptualization and design: S.K., M.C., S.L.W., P.C., R.B.; data collection: S.K., M.C., S.L.W., M.W-M., P.C.; analysis: S.K., M.C., S.L.W., M.W-M., P.C., preparation of the manuscript: S.K., M.C., S.L.W, M.W-M, P.C., R.B.

