## Supplementary data for "SARS-CoV-2 structure and replication characterized by *in situ* cryo-electron tomography"

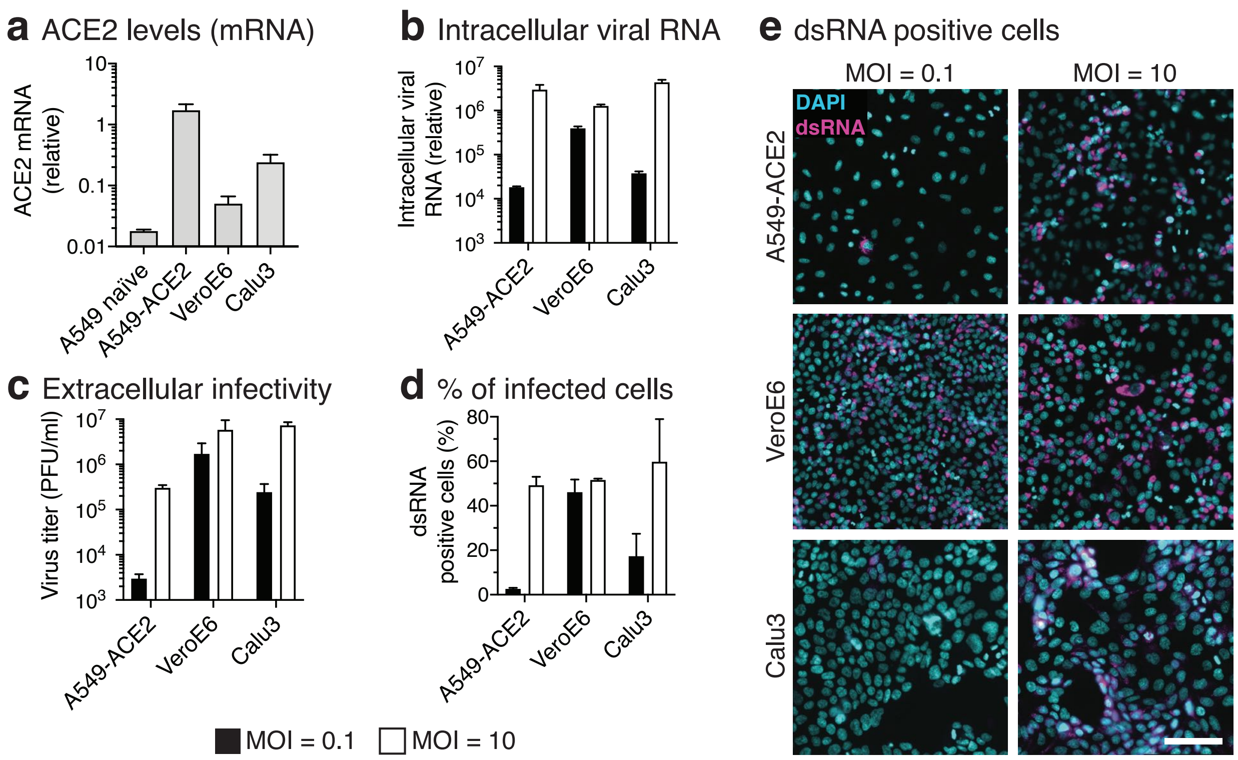


**Figure S1 | ACE2 mRNA levels and SARS-CoV-2 infection in A549-ACE2, VeroE6 and Calu3 cells.** All experiments were performed in triplicates. Data are shown as a bar plot with indicated standard deviation. **a**, ACE2 mRNA levels normalized to HPRT mRNA levels of the three cell lines used in this study determined by RT-qPCR. **b**, Intracellular viral RNA levels (normalized to HPRT mRNA) of the three cell lines infected with SARS-CoV-2 at 16hpi with MOI 0.1 (black) or 10 (white) determined by RT-qPCR. **c**, Released extracellular virus measured as plaque forming units (PFU) per ml for the three cell lines infected with SARS-CoV-2 at 16hpi with MOI 0.1 (black) or 10 (white). **d**, Percentages of dsRNA positive cells for the three cell lines infected with SARS-CoV-2 at 16 hpi with MOI 0.1 (black) or 10 (white). **e**, Wide-field fluorescent light microscopy images used for analysis in (d) showing DAPI stained cell nuclei (cyan) and staining with an anti-dsRNA antibody (magenta) in the three cell lines infected with SARS-CoV-2 16 hpi with MOI 0.1 and 10. Scale bar: 100 µm.

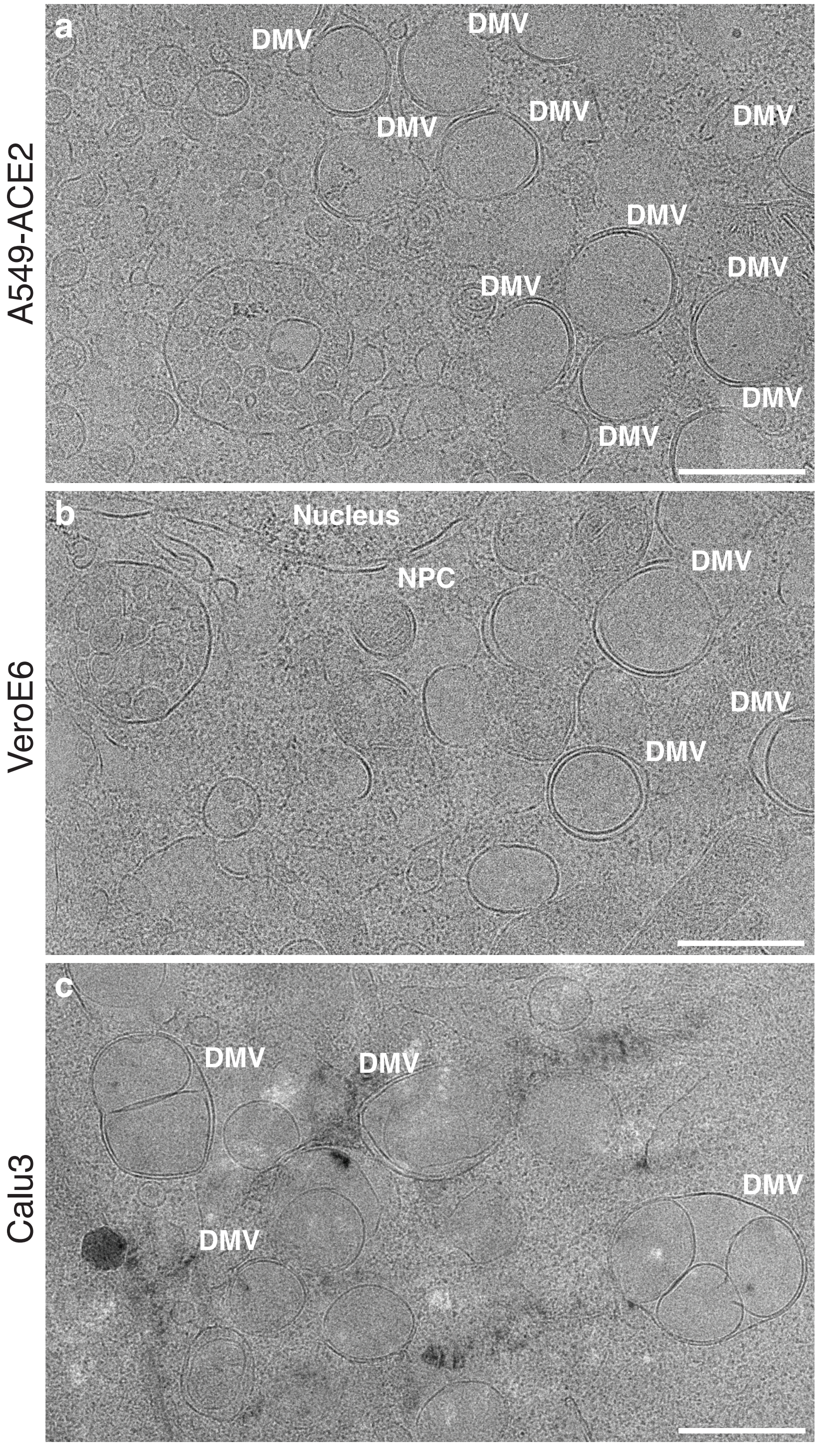


**Figure S2 | Cryo-EM of DMVs in A549-ACE2, VeroE6 and Calu3 cells.** Exemplary images from cryo-EM montages on milled A549-ACE2 (**a**), VeroE6 (**b**) and Calu3 (**c**) cells that were used for tomogram acquisition of DMVs and fused DMVs in Calu3 cells. Due to the thickness of Calu3 and increased clustering of cells, Calu3 cryo-lamellae samples showed partially vitrified regions more frequently. Scale bars: (a–c) 500 nm.

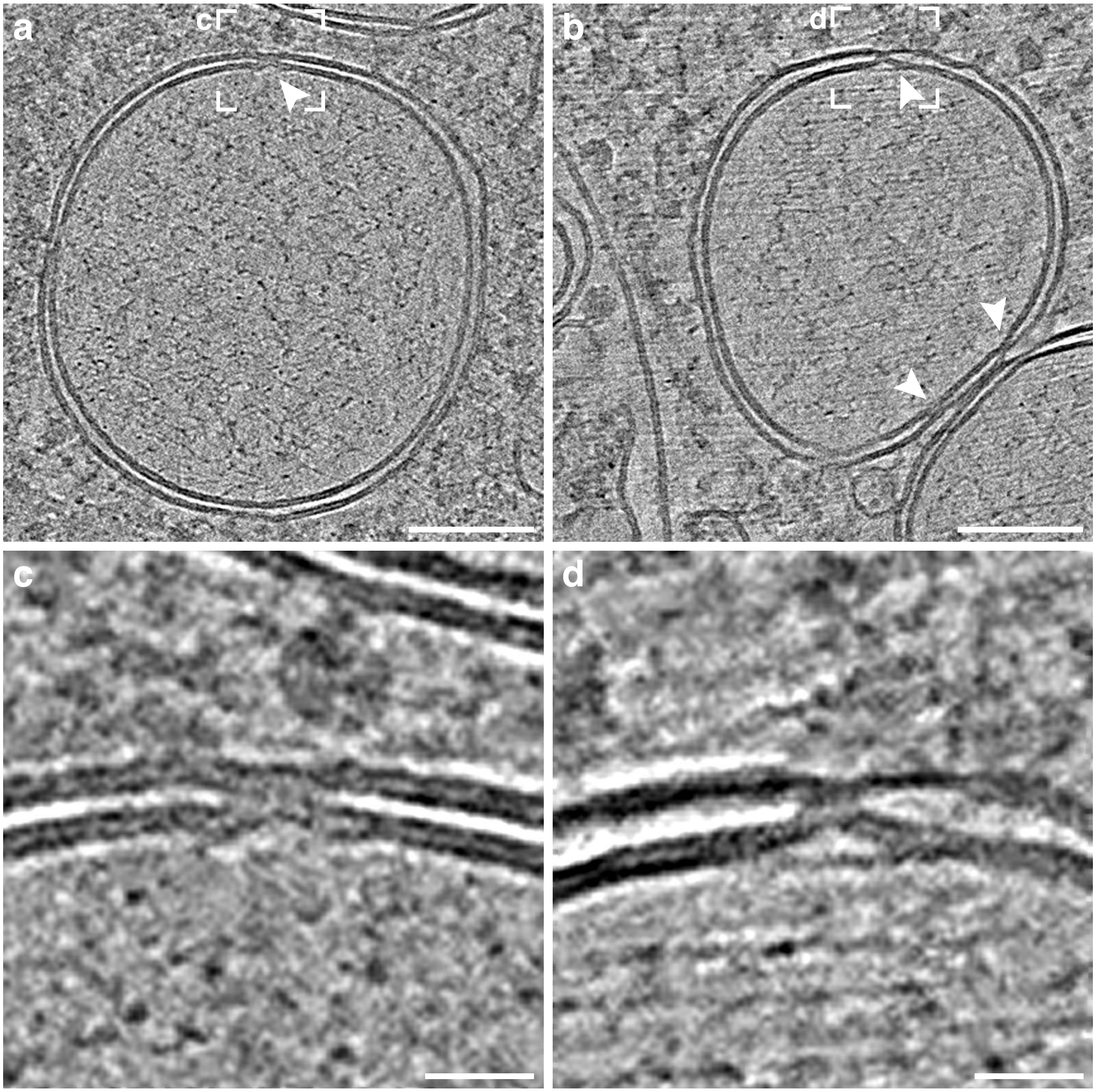


**Figure S3 | Putative pore complexes clamping the inner and outer DMV membrane and connecting the DMV lumen with the cytoplasm. a, b** Slices of tomograms showing DMVs with sites of putative pores (arrowheads), spanning both membranes of the DMV. **c, d**, Magnified images of the putative pore complex. Scale bars: (a, b) 100 nm, (c, d) 20nm

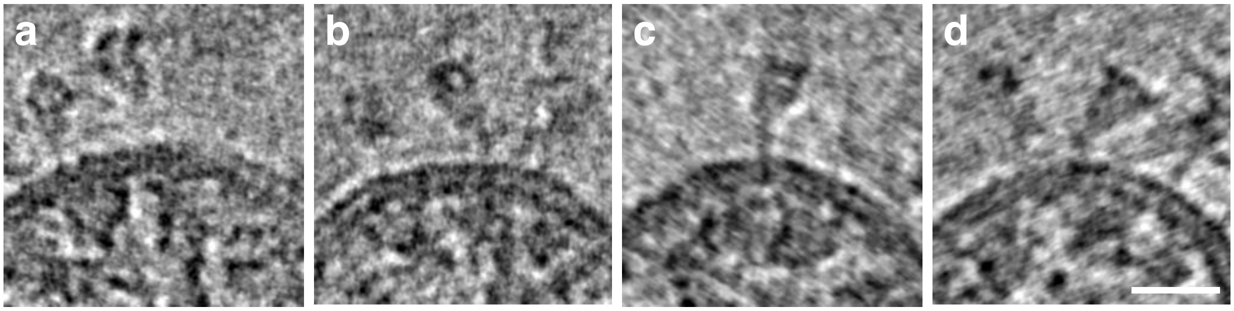


**Figure S4 | Tilted spike glycoproteins of intracellular virions.** Slices of tomograms showing individual S trimers with different tilting angles relative to the viral envelope of 31° (a), 35° (b), 37° (c) and 41° (d). 20 tomogram slices were average and low pass filtered (2 pixel). Scale bar: 20 nm.

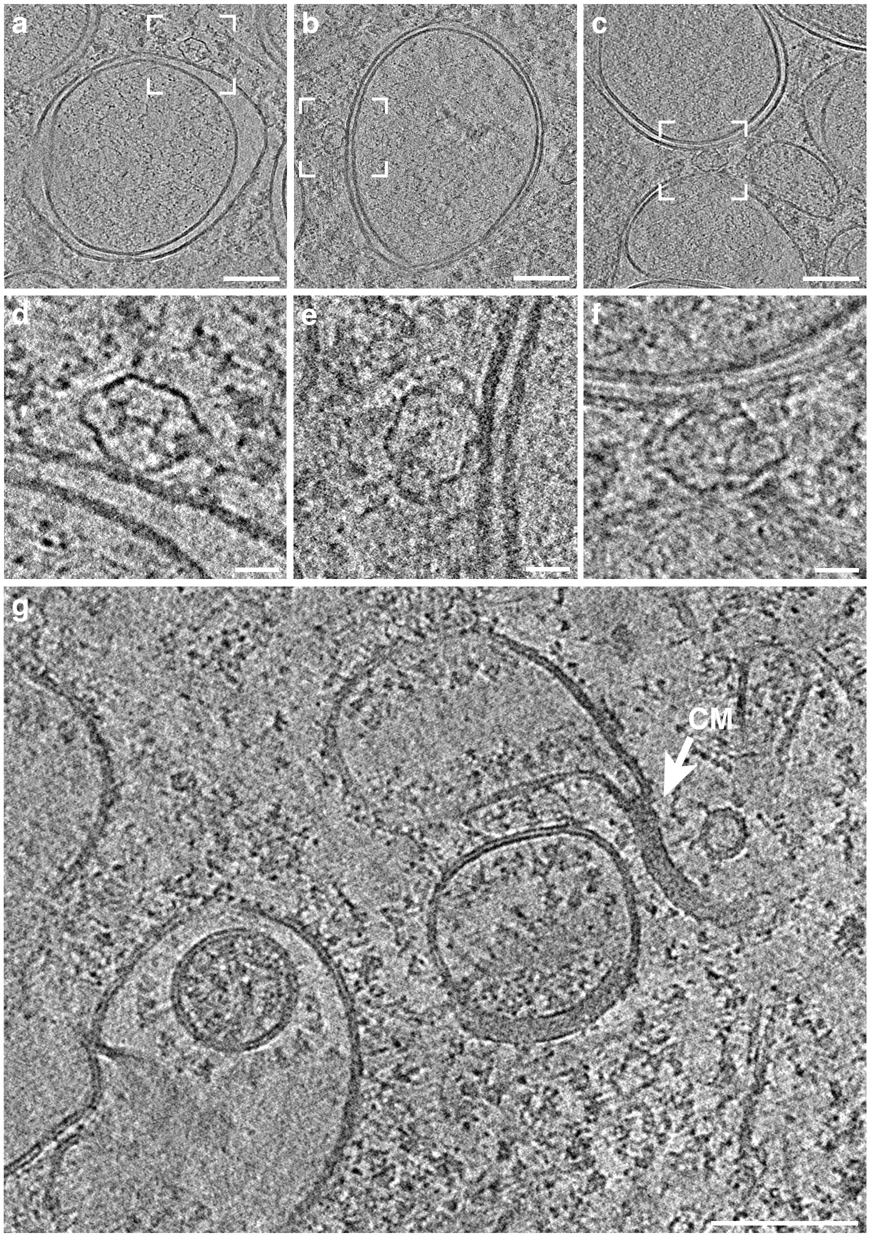


**Figure S5 | Vault protein complex in the proximity of DMV and putative convolute membrane. a–f**, Slices of tomograms (average of 10 slices) showing 3 examples of vault complexes in the proximity of the DMV outer membrane found in SARS-CoV-2 infected VeroE6 (a, c, d, f) and in A549-ACE2 cells (b, e). vault complexes are oriented with the longitudinal axis along the outer membrane in all observed cases. In one case a vault complex was found to be located between two DMVs (c, f). **g**, Slice of a tomogram of SARS-CoV-2 infected Calu3 cell showing a putative zippered ER cisterna (CM; convoluted membrane) characteristic of a density found inside the cisternae. The diameter of the CM is approximately 18 nm. Scale bars: (a, b, c, d, g) 100 nm, (d, e, f) 20 nm.

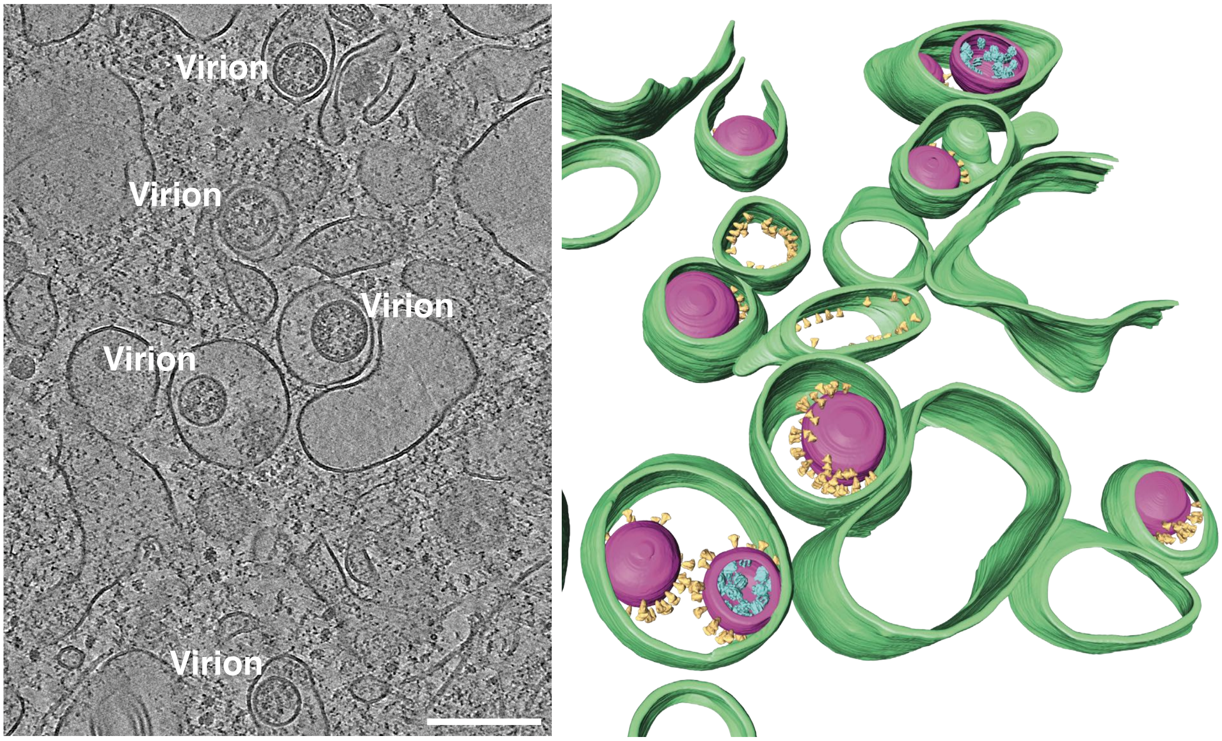


**Figure S6 | Assembled intracellular virions.** Tomogram showing a VeroE6 cell infected with SARS-CoV-2 at 16hpi. 20 slices of the tomogram were averaged, and a non-local means filter was applied. Intracellular released virions inside the ERGIC lumen are indicated. A 3D volume rendering is shown with cellular membranes (green) and viral membranes (magenta). For each S glycoprotein (yellow) and vRNP (cyan) present in the tomogram the subtomogram average of S glycoprotein was placed in the volume rendering. The S trimer average orientations represent the data in the tomogram, whereas the vRNP average orientations were randomized (Movie S3). Scale bar: 200 nm.

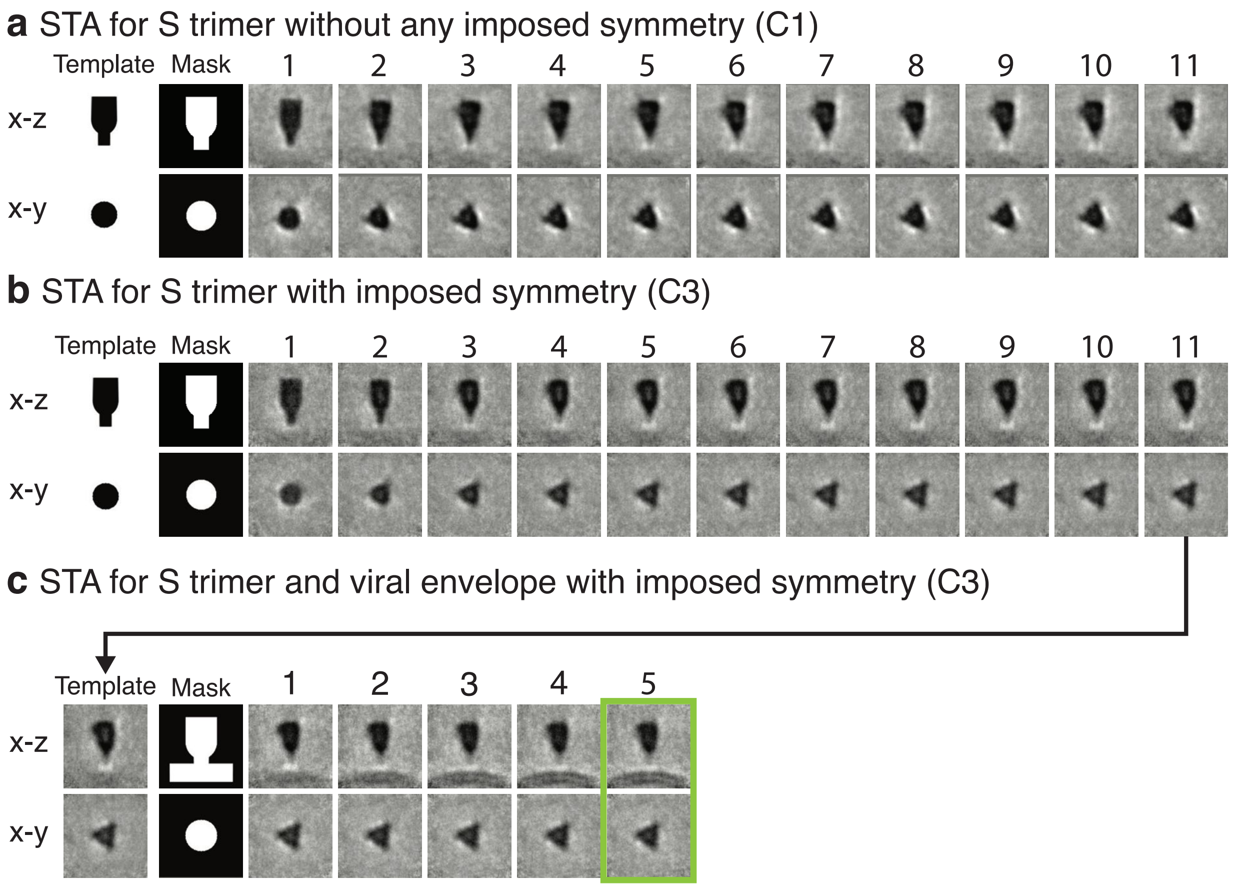


**Figure S7 | Overview of S trimer subtomogram averaging.** Particles (n = 219) corresponding to positions of S trimers in tomograms acquired on cryo-lamella of SARS-CoV-2 infected VeroE6 cells were cropped (160 x 160 x 160 voxels) in Dynamo using the 'general' model. **a**, Particles were iteratively aligned and averaged using a template and mask shown on the left without imposing any symmetry. **b**, Particles were iteratively aligned and averaged using a template and mask shown on the left with c3 symmetry imposed. **c**, Particles were iteratively aligned and averaged using the final average obtained in (b) as a template and a mask shown on the left with c3 symmetry imposed. The final average in B and the final average (green) were c3-symmetrized and used for model fitting in ChimeraX (Movie S4).

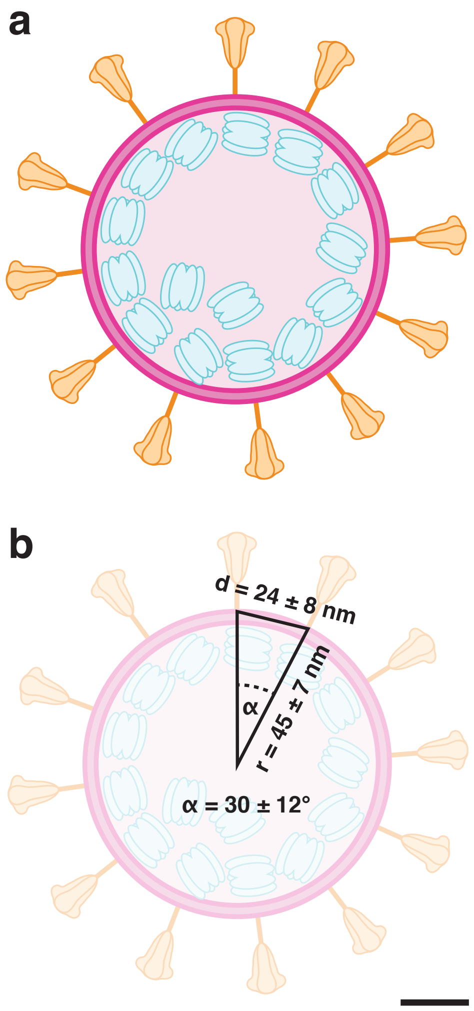


**c** Central angle

$$\alpha = cos^{-1}\left(1 - \frac{d^2}{2r^2}\right)$$

Error central angle

$$\alpha_{err} = \sqrt{\frac{1}{r^2}\left(1 - \frac{d^2}{4r^2}\right)^{-1} d_{err}^2 + \frac{d^2}{r^4}\left(1 - \frac{d^2}{4r^2}\right)^{-1} r_{err}^2}$$

**d**

| Central angle | Number of points | |
|---|---|---|
| 43.71 | 23 | |
| 43.69 | 24 | |
| **41.63** | **25** | $\bar{\alpha} + SD = 42.1°$ |
| 41.04 | 26 | |
| 40.68 | 27 | |
| 39.36 | 28 | |
| 38.71 | 29 | |
| 38.60 | 30 | |
| (…) | | |
| 32.09 | 43 | |
| 31.98 | 44 | |
| 31.32 | 45 | |
| 30.96 | 46 | |
| 30.78 | 47 | |
| **30.76** | **48** | $\bar{\alpha} = 30.4°$ |
| 29.92 | 49 | |
| 29.75 | 50 | |
| 29.37 | 51 | |
| 29.19 | 52 | |
| 28.81 | 53 | |
| (…) | | |
| 19.07 | 122 | |
| 19.01 | 123 | |
| 18.95 | 124 | |
| 18.84 | 125 | |
| 18.78 | 126 | |
| **18.69** | **127** | $\bar{\alpha} - SD = 18.7°$ |
| 18.63 | 128 | |
| 18.56 | 129 | |

**Figure S8 | Model of SARS-CoV-2 with estimation of S trimers per virion. a, b**, Schematic model of a SARS-CoV-2 virion showing the viral envelope in magenta, vRNP in cyan and S trimers in yellow drawn to scale. The two structural transmembrane proteins M and E are not shown. The average radius r = 45 ± 7 nm, average minimum distance between nearest-neighboring S trimers d = 24 ± 8 nm and the corresponding average central angle α = 30 ± 12° are indicated in (b). **c**, Equations used for calculations of the central angle α and its error propagation. **d**, Table of estimates of the 'Tammes Problem' for different numbers of points on a sphere and the corresponding central angle (*14*). The average central angle of 30.4° corresponds closest to a conformation with 48 points on a sphere. For the limits of the confidence interval for the central angle of 18.7° and 42.1° (highlighted in bold), closest matching conformations are 25 and 127 points on a sphere, respectively. Scale bar: 20 nm.

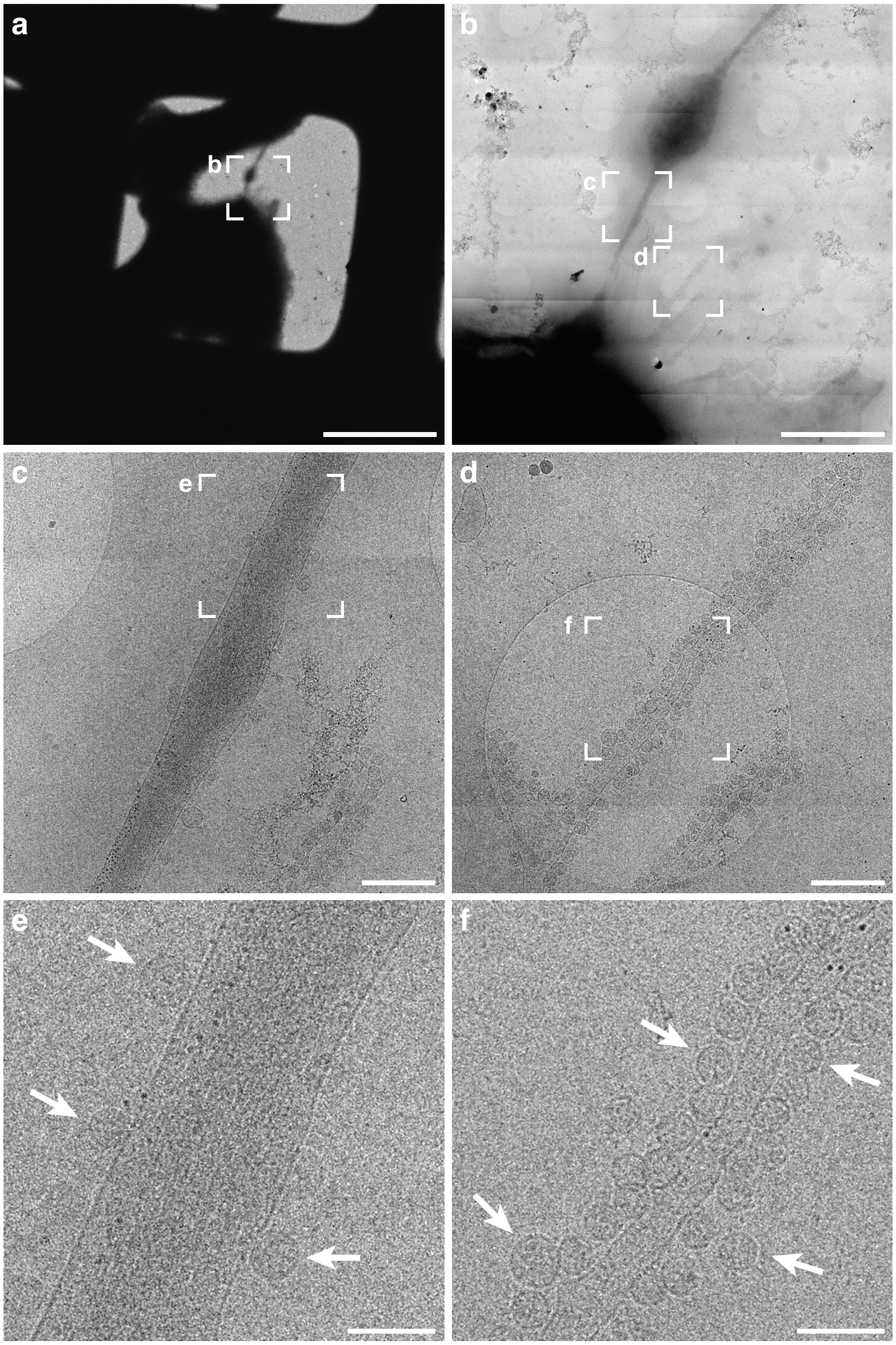


**Figure S9 | Cryo-TEM of extracellular virions attached on VeroE6 cell protrusions.** **a**, Low magnification cryo-TEM image of a 200 Mesh Ti grid coated with a holey carbon (Quantifoil R2/2) with infected VeroE6 cells located in a grid square. **b**, Magnified image of cell periphery denoted in (a) by a dashed square. **c, d**, Magnified images of cell filopodia decorated with extracellular virions denoted in (b) by dashed squares. **e, f**, Magnified images with individual virions (arrows) attached to the filopodia denoted in (c, d) by dashed squares. Scale bars: (a) 50 µm, (b) 5 µm, (c, d) 500 nm, (e, f) 200 nm.

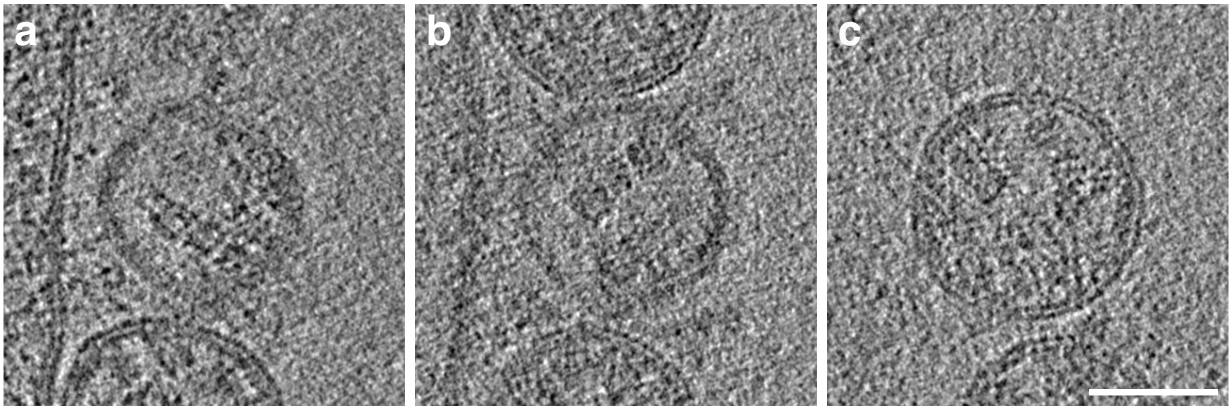


**Figure S10 | Organization of vRNPs in released extracellular virions. a–c,** Slices of tomograms of SARS-CoV-2 virions released from VeroE6 cells. Exemplarily, virions displaying linearly arranged vRNPs are shown from 3 tomograms. For better visualization, 6 slices were averaged from 3x binned and low pass filtered (2 pixel) tomograms. Scale bar: 50 nm.

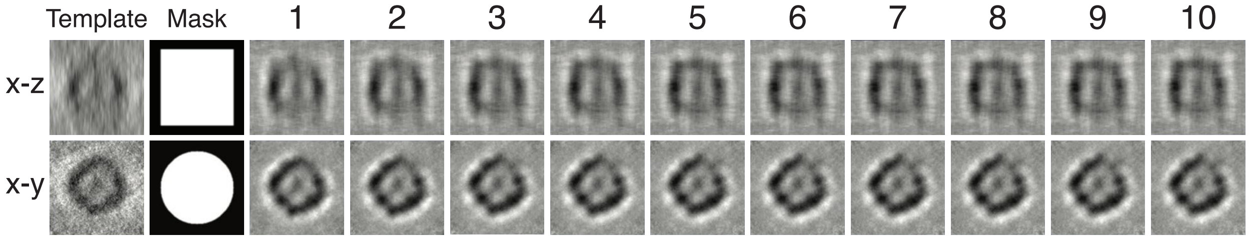


**Figure S11 | Overview of vRNP subtomogram averaging.** Particles (n = 1570) corresponding to positions of vRNPs in tomograms acquired on cryo-lamella of SARS-CoV-2 infected VeroE6 cells were cropped (168 x 168 x 168 voxels) in Dynamo using the 'general' model (*7*). 200 of randomly oriented particles were averaged to generate an initial template. Subsequently, particles were iteratively aligned and averaged using the template and a cylindrical mask (Movie S5).

**On-lamella cryo-ET**

| Infection and plunging | Cells | MOI | Infection hpi | Number of lamellae | Number of tomograms | Number of tomograms used for analysis |
|---|---|---|---|---|---|---|
| Sample preparation I | A459-ACE2 | 7 | 16 | 6 | 11 | 7 |
| | A459-ACE2 | 20 | 16 | 6 | 10 | 2 |
| | A459-ACE2 | 7 | 16 | 7 | 18 | 7 |
| Sample preparation II | Calu3 | 5 | 16 | 12 | 7 | 3 |
| | VeroE6 | 0.1 | 16 | 8 | 4 | 4 |
| | VeroE6 | 0.1 | 16 | 6 | 8 | 4 |
| | VeroE6 | 0.1 | 16 | 8 | 8 | 2 |
| | A459-ACE2 | 20 | 16 | 7 | 2 | 1 |
| **Total** | | | | **60** | **68** | **30** |

**Whole-cell cryo-ET**

| Infection and plunging | Cells | MOI | Infection hpi | Number of tomograms | Number of tomograms used for analysis |
|---|---|---|---|---|---|
| Sample preparation I | A459-ACE2 | 20 | 16 | 5 | 0 |
| Sample preparation II | A459-ACE2 | 20 | 16 | 3 | 2 |
| | VeroE6 | 0.1 | 16 | 14 | 11 |
| | Calu3 | 5 | 16 | 5 | 2 |
| | VeroE6 | 0.1 | 16 | 13 | 6 |
| **Total** | | | | **40** | **21** |

**Table S1 | Sample preparation overview.** Summary of biological replicas, number of milled lamellas, number of acquired and analyzed tomograms from each cell line used in the study.

**Movie S1 | Tomogram of DMV with volume rendering and vRNA segmentation.** 3D rendering of one tomogram (Fig. 1) of A549-ACE2 cells infected with SARS-CoV-2 at 16 hpi showing a DMV's outer and inner membrane (dark and light green, respectively) as well as manually segmented filaments in the DMV's core, color-coded by total filament length. The tomogram was denoised using cryo-CARE.

**Movies S2 and S3 | 3D rendering of SARS-CoV-2 virion budding and assembly at the ERGIC membrane.** 3D rendering from two tomograms (Fig. 3 and S6) of VeroE6 cells infected with SARS-CoV-2 at 16 hpi showing budding events at the ERGIC membrane and intracellular released virions inside the ERGIC lumen are indicated. Cellular and viral membranes are shown in green and magenta, respectively. S trimers (yellow) and vRNPs (cyan) are represented as subtomogram averages. The S and vRNP locations correspond to the location in the tomogram, vRNP orientations were randomized. A non-local means filter was applied on the tomogram, and 20 slices were averaged.

**Movie S4 | 3D rendering of subtomogram average of S trimer with fitted structures.** S trimer subtomogram average (yellow) and the virion envelope (magenta). Fitted structure of the S trimer ectodomain (PDB:6VXX) and the HR2 domain (PDB:2FXP) are shown in black and orange, respectively.

**Movie S5 | 3D rendering of subtomogram average of vRNP.** vRNP complex rotated around the long and short axis.
